# The defence island repertoire of the *Escherichia coli* pan-genome

**DOI:** 10.1101/2022.06.09.495481

**Authors:** Dina Hochhauser, Adi Millman, Rotem Sorek

## Abstract

In recent years it has become clear that anti-phage defence systems cluster non-randomly within bacterial genomes in so-called “defence islands”. Despite serving as a valuable tool for the discovery of novel defence systems, the nature and distribution of defence islands themselves remain poorly understood. In this study, we comprehensively mapped the repertoire of defence islands within >1,300 strains of *Escherichia coli,* the most widely studied organism in terms of phage-bacteria interactions. We found that defence islands preferentially integrate at several dozens of dedicated integration hotspots in the *E. coli* genome. Defence islands are usually carried on mobile genetic elements including prophages, integrative conjugative elements and transposons, as well as on other genetic elements whose nature of mobilisation is unclear. Each type of mobile genetic element has a preferred integration position but can carry a diverse variety of defensive cargo. On average, an *E. coli* genome has 4.5 genomic hotspots occupied by a defence system-containing mobile element, with some strains possessing up to eight defensively occupied hotspots. Our data show that the overwhelming majority of the *E. coli* pan-immune system is carried on mobile genetic elements that integrate at a discrete set of genomic hotspots, and explains why the immune repertoire substantially varies between different strains of the same species.

## Introduction

Bacteria are engaged in a continuous arms race in which they have evolved to defend themselves against the expanding arsenal of weapons at the disposal of phages (1). To this end, they possess dedicated defence systems that protect against phage infection through a variety of molecular mechanisms (2, 3). Many defence systems used by bacteria were only discovered in the past few years, and it is estimated that many additional anti-phage mechanisms are yet to be discovered (4–8).

Bacterial anti-phage defence systems were shown to be non-randomly distributed in microbial genomes (6, 9, 10). Such systems were observed to frequently co-localise in bacterial and archaeal genomes, forming so called “defence islands”: genomic regions in which multiple defence systems cluster together (6, 9, 10). The tendency of defence genes to reside next to one another has enabled the discovery of dozens of novel phage resistance systems based on their genomic presence next to known defence systems (4, 6, 7, 11–15).

Although defence islands have served as a remarkably useful tool for the discovery of new defence systems, reasons for the genomic co-localisation of defence systems and the nature of defence islands themselves remain poorly understood. Recent evidence suggests that defence islands are frequently carried on mobile genetic elements (MGEs). These include integrative and conjugative elements (ICEs) (16), transposons (17), and prophages (5, 8). It was shown that these MGEs can possess dedicated hotspots for carrying multiple anti-phage defence systems. Furthermore, several independent studies have demonstrated that MGEs carrying defence islands were directly responsible for differential phage resistance profiles in closely related strains of *Vibrio cholera* and *V. lentus* (16, 18). It has been hypothesised that anti-phage defence systems carried by MGEs participate in inter-MGE warfare and play a role not only in defending the host bacterium against invading phages, but also in protecting resident MGEs against invading MGEs (19).

In the current study we set out to map and study the repertoire of mobile defence islands in the *Escherichia coli* pan-genome. *E. coli* is the most well characterised model organism for bacteria-phage interactions, but the arsenal of defence islands in its genome and their preferred mode of mobilisation have never been studied thoroughly. By analysing over 1,300 *E. coli* genomes, we demonstrate that MGEs carrying defence islands have a marked preference of defensive cargo, as well as preferred integration hotspots within the *E. coli* genome. Our analysis forms a repository of defence islands in *E. coli* strains, a database that may serve as a resource for the discovery of new defence systems in the future.

## Results

In order to find hotspots for integration of defence system-containing mobile elements in the *E. coli* pan-genome, we examined 1,351 *E. coli* genomes downloaded from the IMG database (20). Each genome was scanned for regions containing genes known to be involved in anti-phage defence, searching for such regions present in some genomes but missing from others (Methods, Figure 1A). We then mapped these regions, henceforth called defence islands, to the reference genome of *E. coli* strain K-12 MG1655, a commonly used laboratory strain whose genome is well characterised. The defence islands mapped to 41 discrete hotspots, most of them empty (unoccupied) in the reference K-12 genome (Figure S1; Table S1).

**Figure 1.**
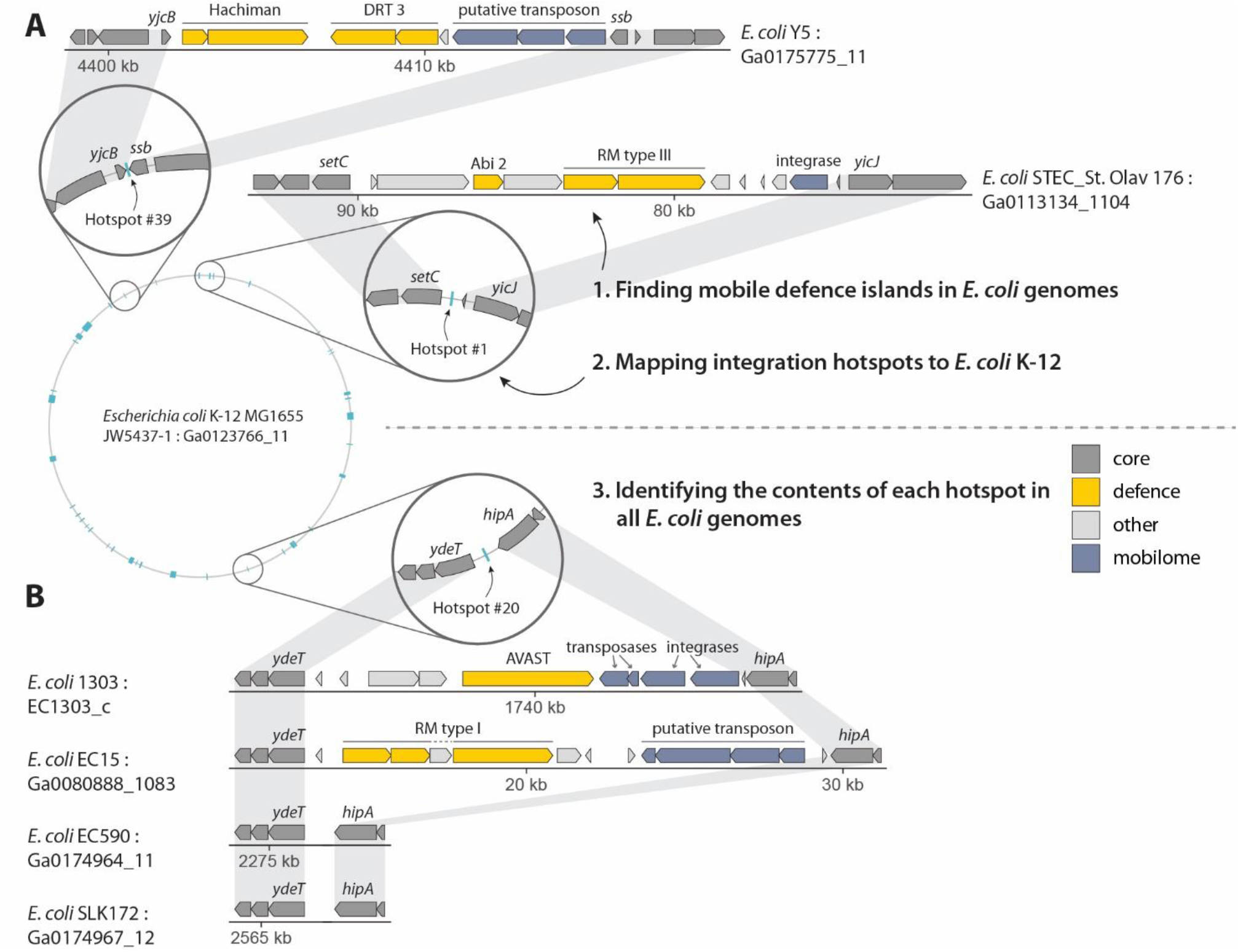
Schematic of the defence island search approach employed in this study. The accession number of each genomic scaffold in the IMG database (20) is shown. Grey shading indicates gene conservation. Genes involved in defence are marked in yellow; Hachiman is a defence system described in (6); DRT and AVAST are described in (4). (A) Regions containing defence systems in 1,351 *E. coli* genomes are mapped to the *E. coli* K-12 genome based on flanking core genes, identifying hotspots for integration of defence-carrying mobile genetic elements. (B) Each hotspot is then searched for in all other *E. coli* genomes, in order to characterize the hotspot occupancy in the pan-genome.

To understand the occupancy of defence island hotspots in the *E. coli* pan-genome, we used the core genes immediately flanking each hotspot in the *E. coli* K-12 reference genome to map these hotspots in the 1,351 downloaded genomes (Figure 1B). With a few exceptions, a given hotspot was unoccupied in the majority of genomes in which it was detected, with a median of 8% occupancy per hotspot (Figure 2A). An exception was the type I-E CRISPR-Cas locus at hotspot #7, which appears to be part of the core genome of *E. coli* and is not found on a mobile genetic element (21). This locus was present in ~70% of the genomes we analysed, while in the remaining ~30% it was degraded. Another locus that was often occupied was the type I restriction-modification (RM) locus at hotspot #36, which was flanked by a transposable element and occasionally included additional defence systems such as Druantia and type IV RM systems (Figure 2A).

**Figure 2.**
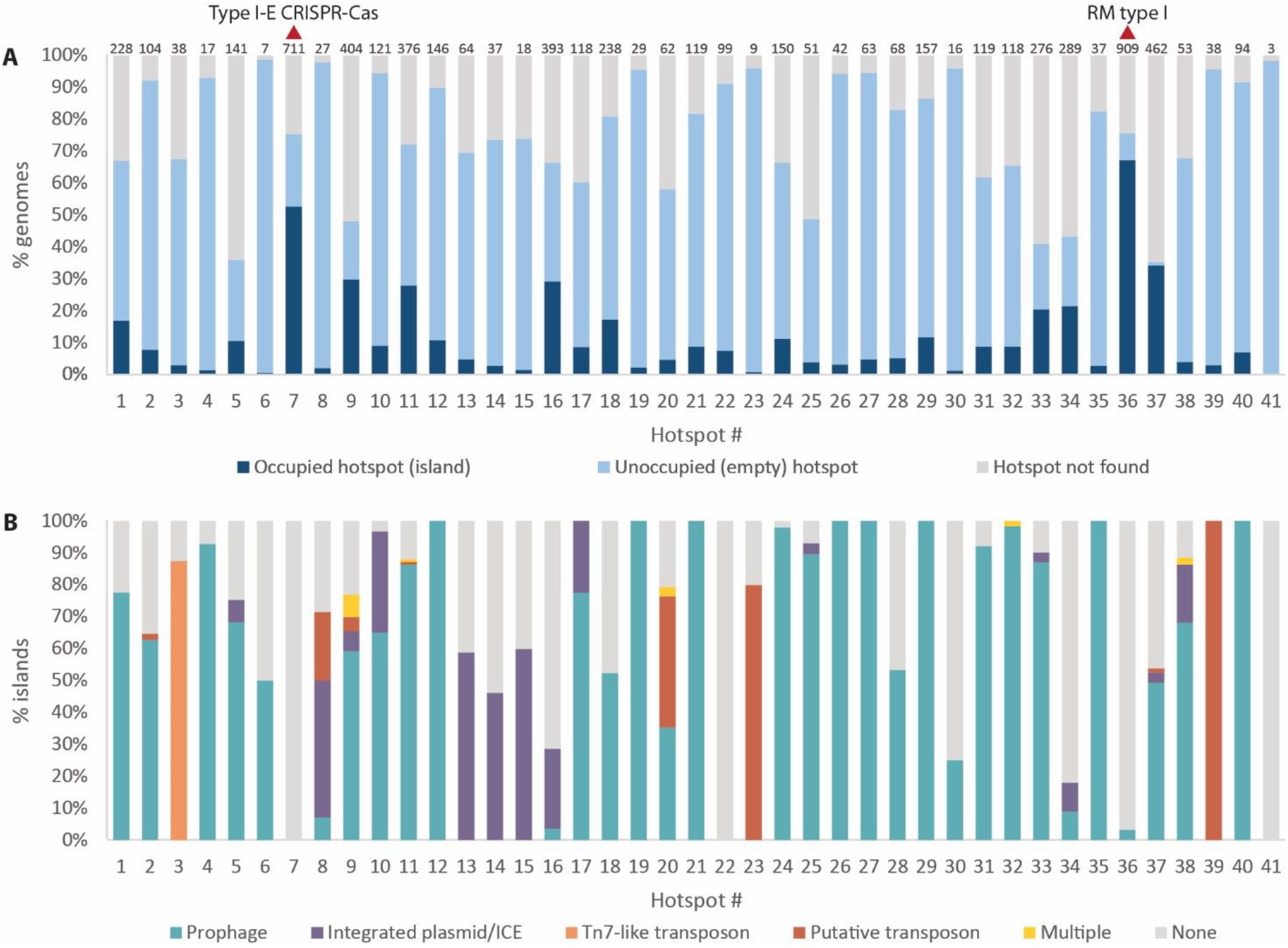
Occupancy of defence island hotspots. (A) Bar graph showing the occupancy of each defence island integration hotspot among 1,351 studied *E. coli* genomes. The number above each bar indicates the number of genomes in which the hotspot was found occupied. (B) Nature of mobile genetic elements integrated at hotspots identified in this study. ICE, integrative conjugative element; multiple, annotations of genes in the integrated element suggest a combination of multiple types of mobile genetic elements.

As expected from a recent analysis of the *E. coli* pan-genome (22), the defence islands we found were mostly carried on mobile genetic elements including prophages and their satellites, transposons, integrative/conjugative elements (ICEs), and integrated plasmids (Figure 2B). Prophages were the most abundant MGE type carrying defence systems (Figure 2B). Mobile genetic elements showed preference for integration at specific hotspots; for example, Tn7-like transposons preferentially integrated at hotspot #3 between the genes *yhiN* and *yhiM,* while P4-like phage satellites that carry defence systems commonly integrated within tRNA genes at hotspots #1, #9, #11, #33, and #37, as well as between genes *yjiA* and *yjiO* at hotspot #36. Non-canonical integrative mobilisable elements including *GISul2* (23, 24) integrated mainly at hotspot #10 and, in some cases, at hotspot #33. These mobile elements do not constitute full conjugative elements but are postulated to hitchhike on other conjugative elements for transfer between species (23, 24).

Some mobile defence islands did not contain any genes identifying them as carried by phage, ICE, transposon, or any other known mobile element. These islands were frequently flanked by genes annotated as “phage integrase” or multiple genes annotated as integrases or recombinases, but no additional phage genes were present in the island (Figure 3A-C). The presence of these islands in only a subset of genomes suggests that they are somehow mobile, but it is not clear how such elements can mobilise between genomes in the absence of known mobility genes. It is possible that multi-gene integrases constitute yet unidentified transposons (Figure 3C), or that integrase genes are remnants of degraded prophages that evolved to become elements dedicated to carrying defence systems, as suggested in (19). It was previously shown that degraded phage elements can parasitize on full-length phages such that during infection by the full phage, some capsids carry the degraded phage genomic material (25). It is possible that some of these integrase-only elements and their cargo are transferred between genomes in a similar manner.

**Figure 3.**
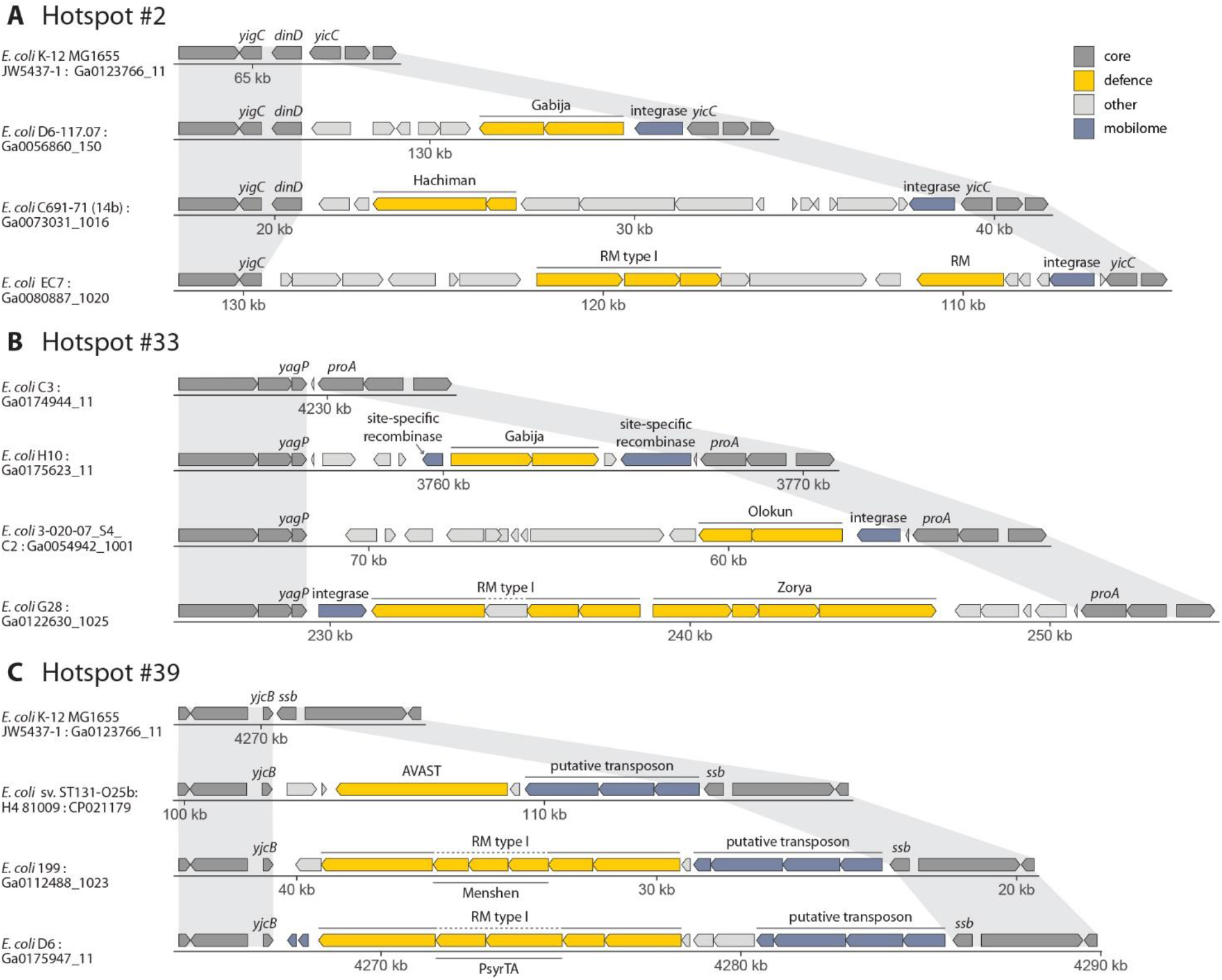
Examples of defence islands with unclear mechanism of mobility. These islands typically contain integrase or recombinase genes but lack other known mobility genes. RM, restriction-modification; Gabija, Hachiman and Zorya are defence systems described in (6); AVAST was described in (4); and Olokun, Menshen, and PsyrTA were described in (7). Gene symbols of flanking core genes are indicated for each hotspot. (A) Selected defence island examples at hotspot #2. (B) Selected defence island examples at hotspot #33. This hotspot is occupied in *E. coli* K-12. (C) Selected defence island examples at hotspot #39.

Overall, we detected 67 types of known defence systems in the hotspots identified in this study (Figure 4). We found that the same hotspot could be occupied by different sets of defence systems in different genomes (Figures 3–5). For example, hotspot #14 could contain CBASS, Hachiman, retron, RM, Septu, and additional systems, often carried on an ICE element that preferentially integrates within a host tRNA (Figure S2; Table S1).

**Figure 4.**
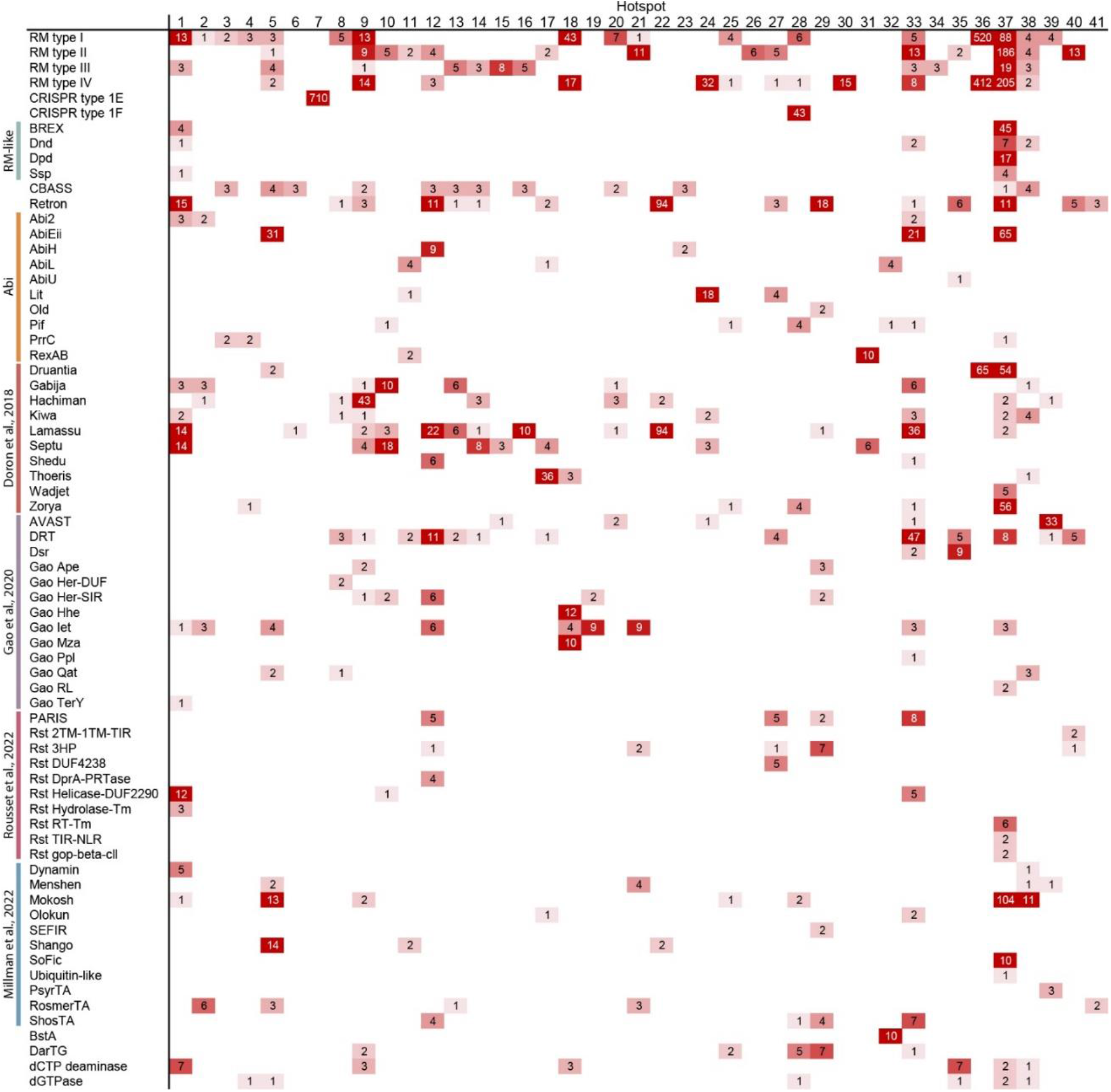
Defence system occupancy in 41 hotspots in the *E. coli* pan-genome. Numbers indicate the occurrences of each defence system within each hotspot.

Some types of defence systems showed preference to be carried by a specific kind of mobile genetic element, or to be integrated at a specific hotspot (Figure 4). For example, the *bstA* gene was found only in prophages integrated at hotspot #32 (Figure 4). BstA is a prophage-encoded abortive infection protein that is naturally silenced by a cognate anti-BstA (*aba*) DNA element and which provides defence against multiple phages that do not carry the *aba* element (26). We found ten instances of BstA-carrying lambda-like prophages, and in all cases these prophages integrated within the tRNA^Arg^ gene at hotspot #32. Similarly, the BREX system (11), which appears in 49 islands in our set, was only present at hotspots #1 and #37; the abortive infection system AbiEii (27) was found only at hotspots #5, #33, and #37; and mobile genetic elements carrying the Wadjet and Zorya defence systems showed preference for integration at hotspot #37 (Figure 4, Figure 5).

**Figure 5.**
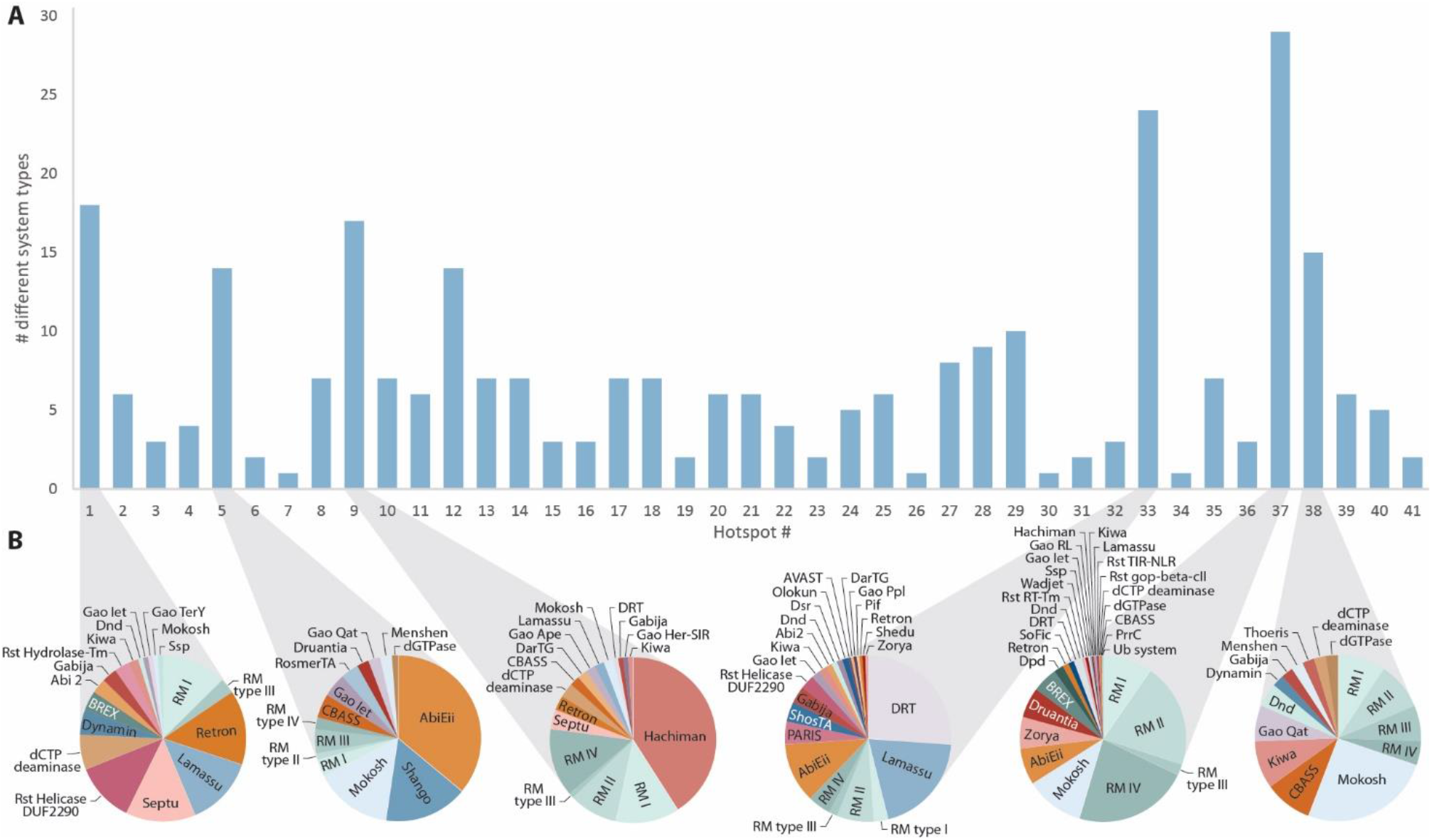
Diversity of defence systems in *E. coli* hotspots. (A) Number of different defence systems found at each of the 41 hotspots mapped in this study. (B) Defence system distribution for a selected set of the most diverse hotspots.

We next examined 190 *E. coli* genomes defined as “finished” in the IMG database (20), i.e., their genomes were completely assembled with no gaps. A given finished genome had, on average, 10.2 of the 41 hotspots occupied with an integrated element, but only a subset of these (between one and eight, 4.5 on average) contained known defence systems (Table S3). Analysing the defence system content of the main chromosome of these genomes using DefenseFinder (28) and PADLOC (29) revealed a total of 1,424 defence systems. Of these, 1,304 (91.6%) were found at the 41 hotspots mapped in the current study (Table S3). Defence systems frequently (53.7% of cases) co-localised with at least one other system on the same island (Table S3), conforming with the previously observed tendency of defence systems to genomically co-localise (6, 9), but also showing that defence systems frequently appear alone (8, 28). Together, these data suggest that the overwhelming majority of the chromosomal defence system repertoire of the *E. coli* pan-genome is carried on mobile genetic elements that preferentially integrate at a discrete set of defined genomic positions.

## Discussion

For many decades, *E. coli* has been the workhorse for studies on interactions between bacteria and phage (30). Many bacterial defence strategies have been discovered, and the mechanisms by which phages evade these defences extensively studied, using *E. coli* as a model organism (30–32). The dataset of mobile defence islands we have collected provides a reference point for defence system occupancy in the pan-genome of *E. coli* and can serve as a resource for future studies aimed at examining the *E. coli*-phage relationship.

Our data show that the vast majority of the defence systems in the *E. coli* pan-genome are carried on mobile genetic elements, explaining the immense variability in presence and absence of these defence systems among different strains of the same species (1). These data are consistent with recent studies showing that in strains of *Vibrio* species, mobile genetic elements carrying defence systems comprise the majority of accessory genes in the pan-genome (16, 18). Indeed, recent studies have detected new defence systems specifically in *E. coli* prophages (8, 26) or have relied on defence-rich hotspots within prophage genomes to reveal new defence systems (5). Future studies will be required to examine whether the same principles of defence system mobility hold true for bacterial species beyond the Proteobacteria phylum.

Many coliphages are strain-specific, infecting only a subset of *E. coli* strains (33). With our map of defence systems localised to *E. coli* defence islands, it will be possible to ask whether mobilisation of defence-containing elements leads to resistance against particular phages, and to potentially identify the specific defence systems preventing phage infection. Moreover, with our accurate mapping of the boundaries of defence islands, it will be possible to search these islands for yet-undiscovered defence systems, thus empowering future studies aimed at expanding the current knowledge of bacterial defence.

Our study, as well as recent studies by others (5, 7, 8, 16, 18, 19), demonstrates that defence systems cluster within specific regions in MGEs, yet the reasons for this clustering are not entirely clear. It is likely that this aggregation of defence systems within mobile elements provides fitness benefits to recipient bacteria living in a phage-rich environment (1). It has also been suggested that synergism between defence systems may promote their co-localisation within and co-transfer between genomes (19, 34). Genes involved in bacterial pathogenicity are frequently mobilised between bacteria on “pathogenicity islands”, which constitute MGEs carrying clusters of pathogenicity genes (35). Similarly, multiple antibiotic resistance genes tend to cluster on plasmids and other mobile elements (36, 37). It is possible that the same evolutionary forces acting to aggregate pathogenicity and antibiotic resistance genes on the same mobile element could act to promote defence system aggregation within defence islands. Understanding the exact nature of these evolutionary forces, as well as the benefits and costs of hosting mobile elements encoding defence islands, awaits future studies.

## Methods

### Defining defence islands in *E. coli* genomes

Prokaryotic genomes were downloaded from the IMG database (20) on October 2017, and all proteins from these genomes were grouped by sequence similarity using MMSeqs (38) to form clusters of homologs, as previously described (7). The subset of *E. coli* genomes in the downloaded dataset was then further analysed. Genome assemblies that were highly fragmented and comprised more than 200 contigs were discarded. The following defence systems were identified in the *E. coli* genome dataset based on cluster annotation, as previously described (7): RM, CRISPR, BREX (11), CBASS (13), DISARM (12), Dnd (39), Druantia, Gabija, Hachiman, Kiwa, Lamassu, Septu, Shedu, Thoeris, Wadjet, Zorya (6), pAgo (40), retrons (14), STK2 (41), Aditi, Azaca, Bunzi, Dazbog, Dodola, Menshen, Olokun, Shango (7), dCTP deaminase (42), DSR1, DSR2, Gao Qat, Gao SIR2-HerA, RADAR (7), PrrC, and Abi proteins (pfams PF07751, PF08843, PF09848, PF10592, PF14253, PF14355).

To determine the boundaries of each putative defence island, the genomic regions upstream and downstream of the systems were scanned until reaching “flanking genes” that are part of the core genome, i.e., belonging to protein clusters found in more than 80% of *E. coli* genomes. Mobile defence islands were defined as defence system-containing regions of at least ten genes which were present in some genomes but absent in others, i.e., their flanking genes were found adjacent to one another in at least one *E. coli* genome.

### Mapping defence islands to the *E. coli* K-12 reference genome

In order to precisely map integration hotspots of mobile defence islands to the *E. coli* K-12 reference genome, clusters of the flanking genes of each defence island were compared to clusters in the genome of *E. coli* K-12 MG1655 JW5437-1 (Genbank accession: CP014348) until a syntenic region was found between the genomes. This was defined as a block of five flanking consecutive genes in the same order and with the same respective clusters as five consecutive genes in the K-12 genome. For cases of gene deletions or duplications in the flanking regions of the defence islands, these hotspots were manually inspected to define the exact integration hotspot and the precise flanking genes (Table S1).

### Using K-12 flanking genes to determine island occupancy in all *E. coli* genomes

Each hotspot was searched for in all analysed *E. coli* genomes using the two genes immediately flanking the hotspot in the K-12 genome (Table S1). To exclude fragmented contigs, only contigs with more than 20 genes were considered. For cases where the immediately flanking genes were not found in the target genome, or in cases where multiple instances of immediately flanking genes were found, a window of ten genes on either side of the flanking genes in K-12 was searched for in the target genome, requiring at least five of the genes matching in gene order and cluster identity. If multiple matches were found, the closest sets of upstream and downstream flanking genes were selected to define the hotspot. Both flanking regions were required in the same contig to declare a hotspot. The resulting islands at these hotspots were filtered for those containing 200 or fewer genes and were defined as ‘empty’ if they comprised three or fewer genes.

### Clustering islands by sequence similarity and manually curating representative islands

In order to remove redundancy in the resulting islands, the nucleotide sequences of the islands at each hotspot were clustered using the cluster module in MMSeqs2 release 12-113e3 (38), with the parameters --cov-mode 0 -c 0.8 --cluster-mode 2 --min-seq-id 0.6 -s 8 --threads 1. The MMSeq2-determined representative sequence was taken from each cluster as a representative sequence of the island. All representative island sequences were manually curated to adjust their flanking genes where necessary (e.g., in the cases of pseudogenised or repeating genes).

### Identifying mobile genetic elements in islands

Islands were inspected for genes associated with mobile genetic elements (MGEs). Type of MGE was determined as described below.

#### Prophages and their satellites

Prophage genes were identified using BLAST search (43) of the amino acid sequences of genes in each island against the mobileOG database (44). The best BLAST hit was taken, if it had at least 80% identity, and 60% coverage of target gene. A gene was classified as prophage if its best hit was of the major mobileOG category ‘phage’, but not of the minor categories ‘infection,regulation’ or ‘chaperone’, which did not appear to constitute phage genes. Additionally, genes were classed as prophage if they had an annotation of ‘capsid’, ‘tail protein’, ‘tail fiber’, ‘tail assembly’, ‘tail sheath’, ‘tail tube’, ‘phage’, but not ‘plasmid’ (excluding genes that were labelled ‘phage or plasmid relaxase’, for example). Clusters of genes homologous to genes from cryptic prophages previously characterised in the *E. coli* genome (45) were also included, as were genes annotated ‘Psu’ and ‘Ogr’, used to classify P4-like phage satellites (46).

#### Integrated plasmids and conjugative elements

Genes from integrated plasmids and conjugative elements were identified if their product annotations included the substrings ‘conjug’, ‘Trb’, ‘type IV secret’, ‘Tra[A-Z]’, ‘mob’, ‘Mob’, ‘Vir[A-Z][0–9]’, ‘t4ss’, ‘plasmid’, but not ‘resolvase’, ‘TraR’ (a transcriptional regulator of phage and bacteria as well as plasmids), ‘TraT’ (also carried by phage), ‘TraW’ and ‘TraU’ (only appeared once in a phage context), ‘ParM’ (the plasmid segregation actin-type ATPase ParM is also found in phage), ‘phage’ (as before, several genes are labelled ‘phage or plasmid relaxase’, for example), and ‘VC0181’ (CBASS component annotated as ‘integrative and conjugative element protein, VC0181 family’).

#### Transposons and putative transposons

Tn7-like transposon genes were classified by searching for genes annotated as ‘TniQ’ and manually verifying the presence of the transposon. Cases of conserved cassettes of multiple genes annotated as ‘phage integrase’ or ‘recombinase’, which did not comprise known transposons or other MGEs, were classified as putative transposons and clusters of their genes were utilised to identify all such putative transposons in islands.

When both integrated plasmid genes and prophage genes were detected in the same island, only the major MGE comprising the most genes was considered. In cases where two types of MGEs were clearly integrated within the same island, the island was recorded as comprising multiple MGE types.

### Identifying defence systems in islands

DefenseFinder (28) and PADLOC (29) were utilised to identify known defence systems in each island. Amino acid sequences of genes in all 1,351 *E. coli* genomes were submitted to DefenseFinder release version 1.1.0 (28). Genes predicted to be part of multiple different defence systems were inspected manually for proper annotation. Amino acid sequences and gff3 files of genes in each island were submitted to the PADLOC web server v1.1.0 (29) with defence systems included in padlocdb v1.4.0. Systems annotated by PADLOC as ‘[system]_other’ were excluded, since these represent partial or separated defence systems. When two overlapping systems of the same type were predicted by the two tools, all constituent genes were considered part of this system.

Regions outside hotspots in the main chromosome in all finished *E. coli* genomes were similarly analysed using DefenseFinder. Defence systems identified outside hotspots were further examined manually to exclude cases in which the hotspot was not detected because of the deletion of flanking core genes on one or both sides of the hotspot.

## Acknowledgements

We thank the Sorek laboratory members for comments on earlier versions of this manuscript. R.S. was supported, in part, by the European Research Council (grant ERC-AdG GA 101018520), Israel Science Foundation (grant ISF 296/21), the Deutsche Forschungsgemeinschaft (SPP 2330, grant 464312965), the Ernest and Bonnie Beutler Research Program of Excellence in Genomic Medicine, the Minerva Foundation with funding from the Federal German Ministry for Education and Research, and the Knell Family Center for Microbiology. D.H. was supported in part by a fellowship from the Israel Ministry of Absorption. A.M. was supported by a fellowship from the Ariane de Rothschild Women Doctoral Program and, in part, by the Israeli Council for Higher Education via the Weizmann Data Science Research Center.

## Supplementary Figures

**Figure S1.**
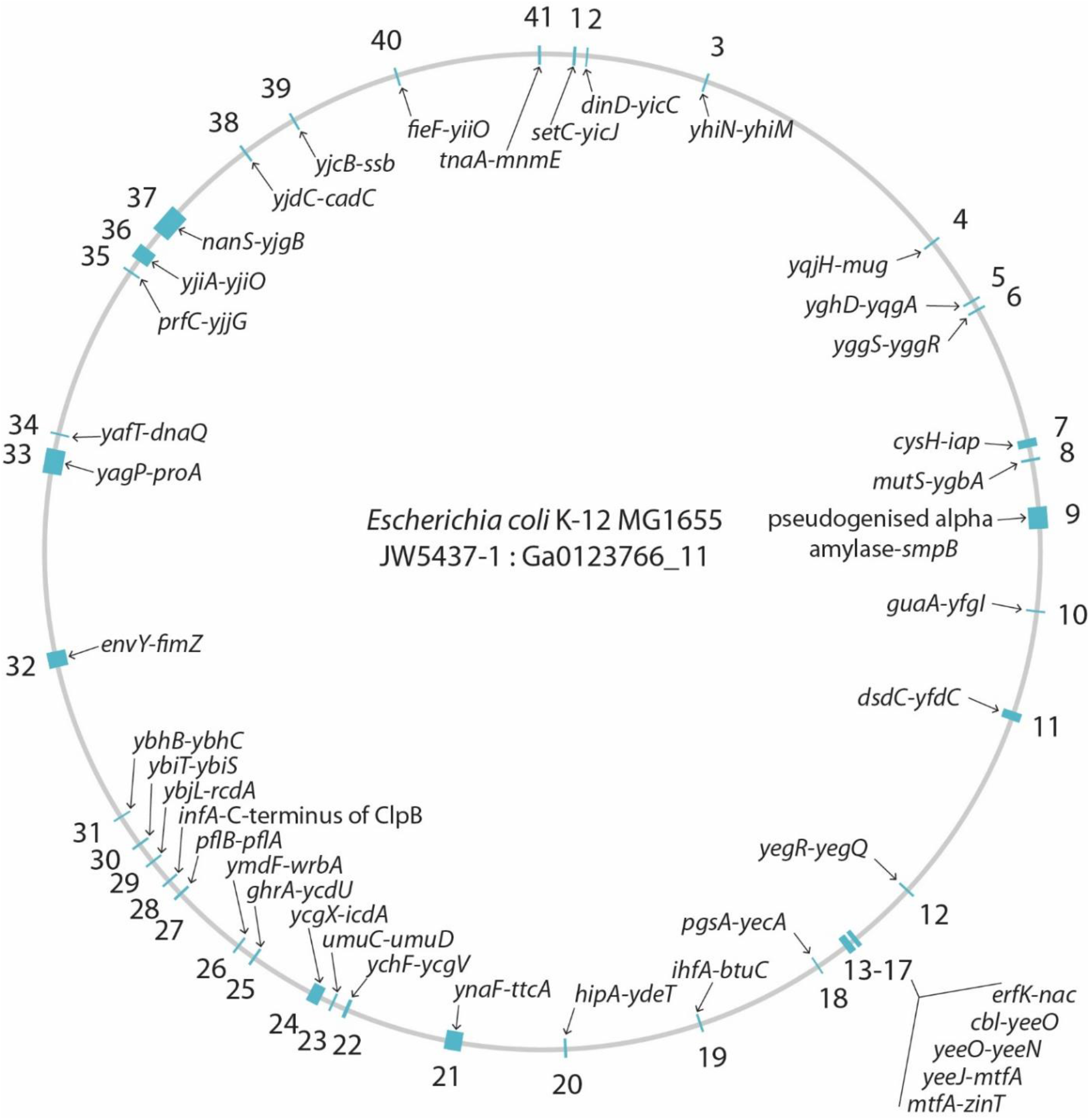
Genomic map of hotspots in the *E. coli* K-12 reference genome. Numbers outside the ring indicate hotspot number. Flanking core genes are indicated for each hotspot on the inside. Blue ticks indicate the position of the hotspot in the E. coli K-12 reference genome, with thicker ticks reflecting hotspots that are occupied in K-12.

**Figure S2.**
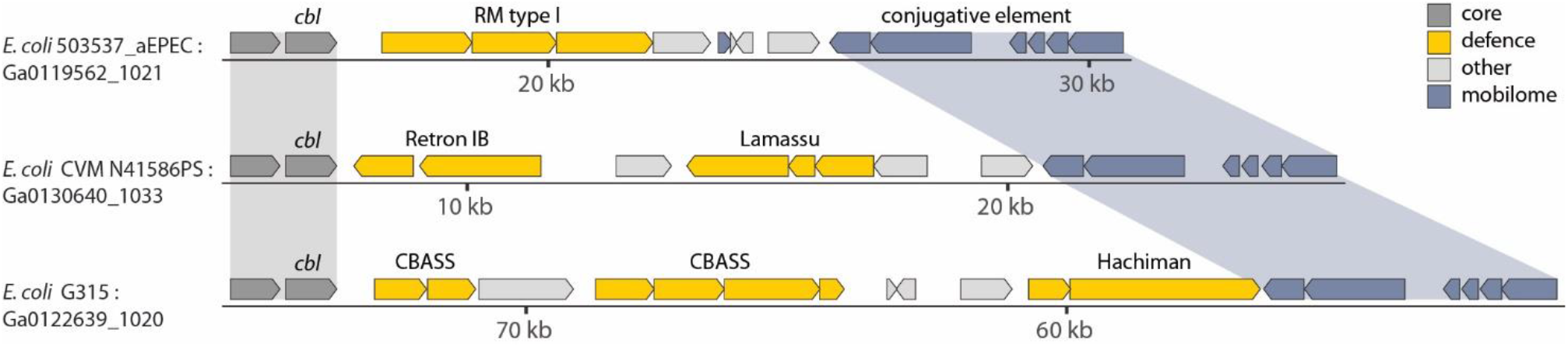
A conjugative element carrying diverse defence systems in a dedicated hotspot. Instances of this element shown (also called “high pathogenicity island” (47)) are all integrated at hotspot #14. Only part of the conjugative element is shown for space constraints.

## Notes

### Competing Interest Statement

R.S. is a scientific cofounder and advisor of BiomX and Ecophage.

